# Chromosome 1 trisomy overcomes the block of sphingolipid biosynthesis caused by aureobasidin A and confers pleiotropic effects on susceptibility to antifungals in *Candida albicans*

**DOI:** 10.1101/2020.04.07.031112

**Authors:** Yi Xu, Feng Yang

## Abstract

Sphingolipids are important membrane lipid components of eukaryotic cells. In *Candida albicans*, chromosome 1 trisomy not only overcame the block of sphingolipid biosynthesis caused by aureobasidin A, but also altered tolerance to three of the four major classes of antifungal drugs. Two haploinsufficient genes on chromosome 1, *PDR16* and *IPT1*, were associated with tolerance to aureobasidin A. This study illustrates an example of multi-drug tolerance caused by aneuploidy in the human fungal pathogen *C. albicans*.

---

*Candida albicans* is a prevalent opportunistic human fungal pathogen (1). Treatment options of candidiasis are very limited, with only four classes of antifungal drugs available: echinocandins, azoles, polyenes and flucytosine (2). More worrisome is the fact that multi-drug tolerance in *C. glabrata* and *C. auris* has been reported (2, 3). Genome plasticity is a hallmark of *C. albicans* (4–7). Aneuploidy, an unbalanced number of chromosomes, widely exists in clinical drug resistant *C. albicans* isolates (8), occurs in the lab under stress conditions (9–11), and is the major mutation during in vivo passages in animals (12, 13). Aneuploidy confers advantage under particular conditions, usually via changing the relative copy number of specific genes because they reside on the aneuploid chromosome (14, 15). Aneuploidy also changes the relative copies of the other genes on the aneuploid chromosome, and their gene products, thus, aneuploidy has the potential to affect multiple phenotypes (16). In *C. albicans*, we have shown that chromosome 5 monosomy causes cross tolerance to caspofungin and flucytosine (10). Chromosome 2 trisomy causes cross tolerance to both caspofungin and hydroxyurea, a chemotherapeutic drug (15).

Sphingolipds are important components of the eukaryotic cell membranes. Sphingolipds influence membrane structure and rigidity (17). Sphingolipids assemble to form dynamic ordered nanoscale microdomains known as lipid rafts, which can act as organizing centers for the assembly of signaling molecules, and segregating of lipids, receptors, adaptors, kinases, scaffolding proteins, and cytoskeletal apparatus (18, 19). Sphingolipids are also important signaling molecules involved in the transport and targeting of membrane proteins (20).

Aureobasidin A (ABA) is a cyclic depsipeptide antifungal antibiotic isolated from the fungus *Aureobasidium pullulans R106* (21). It inhibits the essential inositol phosphorylceramide (IPC) synthase in pathogenic fungi including *Candida* and *Aspergillus* species (22), and is widely used as a fungicidal drug (23). Resistance to ABA is mostly caused by mutations of *AUR1* gene, which encodes the target enzyme of ABA (24, 25). Deletion of several sphingolipid-metabolizing enzymes also causes ABA resistance (26–29).

Using *C. albicans* reference strain SC5314, we did growth curve in YPD broth supplemented with ABA at 37°C. The inoculum was 2×10^3^ cells/ml. The growth was monitored for 24h. We found that ABA inhibited the growth at concentration of 20 ng/ml (Fig. 1A). we then plated 1×10^6^ cells of SC5314 on YPD plates supplemented with 20 ng/ml. The plate was incubated at 37°C for 72h (Fig. 1B). We randomly picked up 18 colonies from the drug plate, and tested the susceptibility to ABA. Only two of the survivors (#1 and #3) were more tolerant to ABA than the parent (Fig. 1C). Whole genome sequencing indicated that the tolerant survivors had duplication of chromosome 1 (Fig. 1D). Interestingly, chromosome 1 trisomy not only caused tolerance to ABA, but also altered tolerance level to unrelated antifungal drugs: increased tolerance to amphotericin B (AMB) and flucytosine (5FC), and decreased tolerance to caspofungin (CSP) (Fig. 2A). Thus, this study is another example of cross tolerance to unrelated drugs mediated by aneuploidy in *C. albicans*. We found two genes on chromosome 1, *PDR16* and *IPT1*, were haploinsufficient. Heterozygous deletion of *PDR16* and *IPT1* in the diploid parent strain was sufficient to cause decreased tolerance to ABA, however, these two genes were not involved in tolerance to AMB, 5FC and CSP (Fig. 2B). Thus, chromosome 1 trisomy alters tolerance to ABA, AMB, 5FC and CSP via regulating different genes. Similarly, we previously found that chromosome 2 trisomy caused cross tolerance to CSP and hydroxyurea, a chemotherapeutic drug, via regulating different genes on chromosome 2 and different pathways (15). In *Saccharomyces cerevisiae*, more copies of *PDR16* was found to confer tolerance to ABA, probably via reducing the effectiveness of ABA against IPC synthase activity (30). In *C. albicans*, screening of a library of 2,868 heterozygous deletion mutants indicated that *IPT1* was haploinsufficient and heterozygous deletion of *IPT1* caused decreased tolerance to ABA (31). This study sheds new light on the mechanism of multi-drug tolerance in *C. albicans*, which is notorious for the genome plasticity.

**Figure 1.**
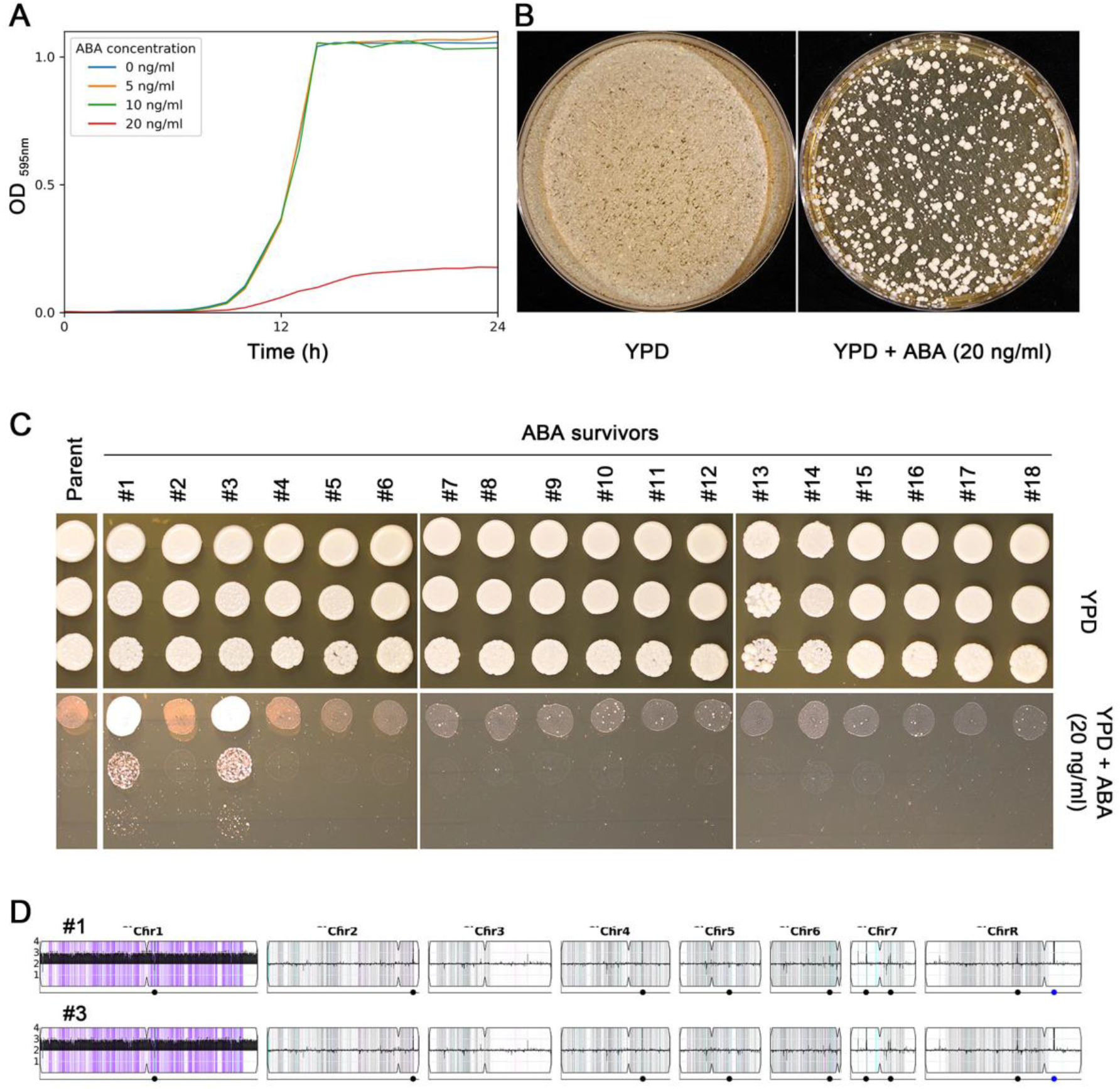
Aneuploidy enables *C. albicans* to survive from inhibitory concentration of aureobasidin A. Measurement of growth curve in YPD broth supplemented with aureobasidin A (ABA) indicated 20ng/ml of ABA inhibited the growth of the reference strain SC5314 (A). On YPD agar plates, some cells could survive from 20ng/ml of ABA (B), but only two (#1 and #3) out eighteen colonies randomly tested obtained tolerance to ABA, as indicated by the spot assay (C). These two strains were deep sequenced and the karyotypes were visualized by using YMAP (D)(32).

**Figure 2.**
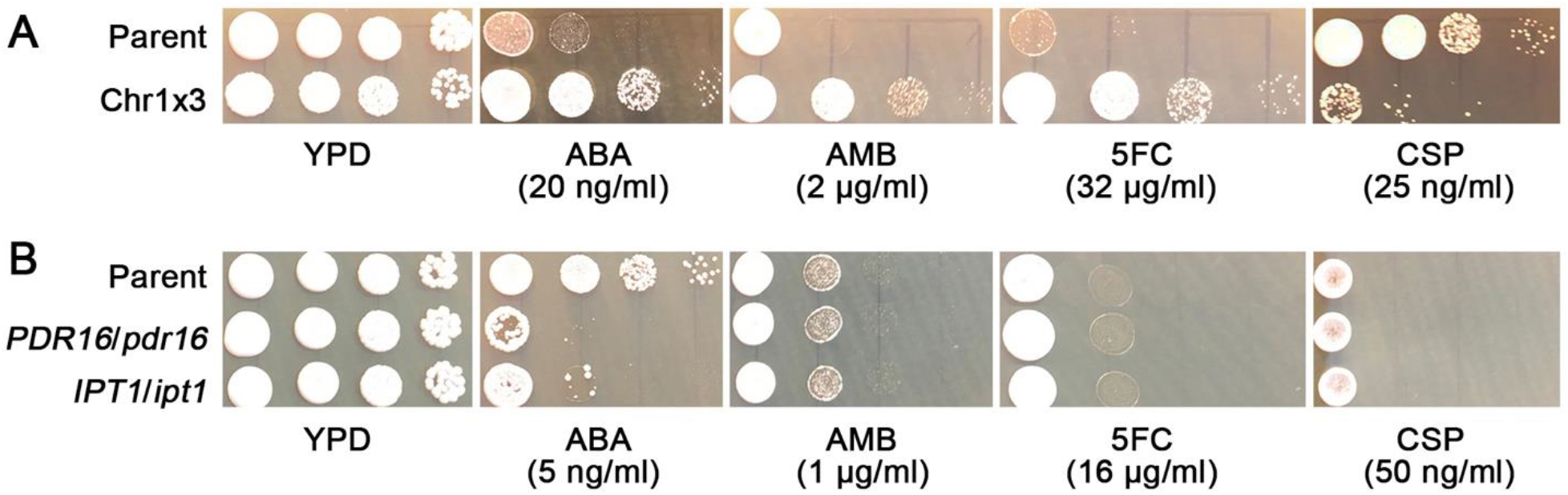
Chromosome 1 trisomy alters tolerance to unrelated drugs. Spot assay indicated that chromosome 1 trisomy conferred increased tolerance to ABA, amphotericin B (AMB), flucotysine (5FC), and decreased tolerance to caspofungin (CSP), as compared to the wild type strain SC5314, which is diploid (A). Two genes on chromosome 1, *PDR16* and *IPT1*, were haploinsufficient. Heterozygous deletion of *PDR16*, or *IPT1*, from the wild type strain, was sufficient to cause decreased tolerance to ABA, but these two genes were not associated with tolerance to AMB, 5FC or CSP (B).

The DNA-Seq data are available in the ArrayExpress database at EMBL-EBI (www.ebi.ac.uk/arrayexpress) under accession number E-MTAB-8942.

## Acknowledgements

This study was supported by the National Natural Science Foundation of China (81402978).

